# Adapter dilution and input optimization for Qiagen QIAseq miRNA Library kit

**DOI:** 10.1101/2025.04.30.651388

**Authors:** Fahri Hasby, Jörg Bachmann, Chuan Wang, Orlando Contreras-Lopez, Anja Mezger

## Abstract

MicroRNAs (miRNAs) are small, non-coding RNA that play a critical role in regulating gene expressions that are important for a multitude of biological processes. In the preparation of miRNA sequencing libraries by using QIAseq miRNA library kit, adapter dimers might occur inadvertently and compromise the sequencing performance. In this technical note, we tested different adapter dilution strategies to minimise or diminish adapter dimers without compromising data quality. In our experiments, we found that RNA input amount plays a larger role in adapter dimer formation rather than the availability of the adapters itself. We suggest preparing the sequencing library with 10 times more RNA than the recommended input amount of the chosen protocol, e.g. following the adapter dilution and PCR cycle number of the standard 1 ng protocol with 10 ng RNA input. We also suggest analysing the data with deduplication by using unique molecular identifiers (UMIs) as it helps remove unusable reads.

## Introduction

MicroRNAs (miRNAs) are small, non-coding RNA molecules, typically 20-24 nucleotides in length that play a critical role in regulating gene expression (Bartel, 2004). By binding to complementary sequences on target messenger RNAs (mRNAs), miRNAs can either degrade the mRNA or repress its translation, thus influencing a wide array of biological processes including development, differentiation, and disease pathology (Lee & Kim, 2005).

Given their extensive regulatory functions, miRNAs are essential for maintaining cellular homeostasis and are involved in numerous physiological and pathological processes. They have been implicated in controlling various cellular activities such as proliferation, apoptosis, and immune responses, making them crucial for understanding normal biology and disease states, including cancer, cardiovascular diseases, and neurological disorders (He *et al*., 205; Calin & Croce, 2006).

Next-generation sequencing is the state-of-the-art method in miRNA quantification. In the preparation of miRNA sequencing libraries, adapter dimers—byproducts of the ligation process where two adapters ligate to each other instead of to the miRNA—can occasionally occur. These adapter dimers can be a significant problem as they consume sequencing reads that would otherwise be available for genuine miRNA sequences (Kim & Kim, 2014). As a result, the presence of adapter dimers can drastically reduce the overall sequencing yield, impacting the depth and quality of the data (Wang & Lin, 2016). To mitigate the negative impact of adapter dimers, it is essential to optimise the concentration of adapters used during library preparation. Proper optimization can reduce the formation of adapter dimers, thereby enhancing the overall quality and quantity of the miRNA sequencing data (Lin & Chen, 2018).

This technical note will discuss how the dilution of adapters, along with the amount of input RNA, can influence key aspects of miRNA sequencing library preparation. These aspects include library yield, the prevalence of adapter dimers, sensitivity, and the overall complexity of the library. By addressing these factors, the note aims to provide practical guidance for optimising miRNA library preparation protocols to achieve high-quality sequencing results.

## Method

### Preparation of sequencing library

The sample used in this experiment is a commercial total RNA that contains miRNA, Universal miRNA Reference Kit (Agilent, USA; 750700). A serial dilution of the original sample was performed with nuclease-free water to achieve 0.2 ng/µl, 2 ng/µl, and 20 ng/µl concentration. The samples were then aliquoted into 100 ng, 10 ng, and 1 ng of total RNA input in 5 µl of nuclease-free water. Each adapter dilution was replicated 7 times and subjected to three different inputs (Table 1). Nuclease-free water was used as a negative control to account for contamination, adapter dimers, and free primers.

**Table 1.** Different dilutions of 3’ and 5’ adapters and input amount used in this experiment.

| Protocol | Input amount | 3' adapter dilution | 5' adapter dilution | RT primer dilution | PCR cycles |
| --- | --- | --- | --- | --- | --- |
| **Standard 10 ng** | 100 ng, 10 ng, 1 ng | 1:5 | 1:2.5 | 1:5 | 19 |
| **Standard 1 ng** | 10 ng, 1 ng | 1:10 | 1:5 | 1:10 | 22 |
| **1 ng 2x dilution** | 10 ng, 1 ng | 1:20 | 1:10 | 1:10 | 22 |

The sequencing library was prepared with QIAseq miRNA Library Automation kit (Qiagen, USA; 331509) and indexing was done with QIAseq miRNA QIAseq miRNA 96 Index IL and IL Auto A (Qiagen, USA; 331565/331569). The QIAseq miRNA Library kit works in an unbiased reaction where adapters are ligated sequentially to the 3’ and 5’ ends of miRNAs. Subsequently, universal cDNA synthesis (with a unique molecular identifier; UMI), cDNA cleanup (bead-based), library amplification, and library cleanup are performed.

RT-PCR primer dilution and PCR cycle numbers were adjusted according to the input amount as instructed in the original protocol from the manufacturer (Table 1). The sequencing library was prepared using a BioMek i7 Hybrid liquid handler (Beckman-Coulter, USA). The library preparation was done in 3 batches, each batch corresponding to a different protocol (Table 1). Negative controls (with nuclease-free water only) were included in all 3 batches. Prior to sequencing, all libraries were subjected to concentration measurements using Qubit dsDNA HS fluorometer (Thermo Fisher Scientific, USA) and fragment analysis using capillary electrophoresis, Fragment Analyzer (Agilent, USA).

### Sequencing setup

All samples, including negative controls, were sequenced on the Illumina NovaSeqXPlus 1.5 B platform, yielding 100 bp single-end sequences (100-8-0-0) with the aim of 265 M clusters (i.e. 5 M clusters per sample). To optimize loading concentration, 80 pM and 160 pM loading concentrations were tested on different lanes.

### Bioinformatic analyses

The entire dataset of raw sequence reads was analysed using nf-core/smrnaseq (v2.3.1). The nf-core/smrnaseq is a bioinformatics best-practice analysis pipeline used for small RNA sequencing data analysis (Peltzer, 2024). The nf-core/smrnaseq pipeline was run with the default settings for QIASeq libraries, or with a manual deduplication process using umi_tools and seqkit (Shen *et al*. 2016, 2024) prior to the analysis using nf-core/smrnaseq pipeline with the same settings. The pipeline uses Nextflow, a bioinformatics workflow tool, and pre-processes raw data from FastQ inputs, aligns the reads to genome reference, and performs extensive quality control on the results. The pipeline generates aligned BAM files and gene count matrices, along with numerous quality control metrics. In this study, the human reference genome (*Homo sapiens, GRCh38*) was used as reference.

## Results & Discussions

### Library preparation and sequencing

In total, 52 sequencing libraries were successfully prepared from all samples except for one replicate of the 10 ng input, prepared with the ‘1 ng 2x diluted’ protocol (Table 1). The lane yield was proportional to theloading concentration– the 160 pM yielded more reads than the 80 pM–and no suboptimal clustering was observed during the sequencing. The average library size from all input and protocol is 184 ± 2 bp, with a median insert size of 22 bp.

Adapter dilution does not affect library yield for low input amount (1 ng). The yield seems to be more affected by the input amount rather than the adapter availability (Figure 1). However, for 10 ng input, higher adapter dilution seems to negatively influence the yield, when comparing the yield on the standard 1 ng protocol and ‘1 ng 2x diluted’ protocol. These two protocols had the same number of PCR cycles, thus the drop in yield can be attributed to the adapter dilution only. The higher number of PCR cycles in the standard 1 ng protocol compared to the standard 10 ng protocol is likely to cause a higher yield in the 10 ng input.

**Figure 1.**
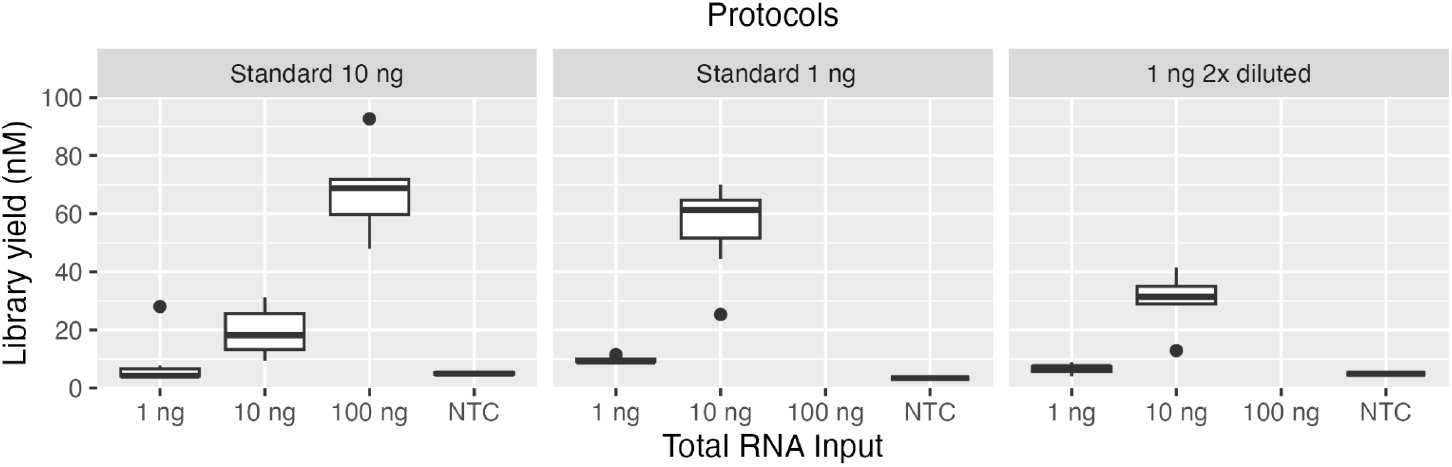
Variations of library yield (nM) across different adapter dilution protocols and inputs. Lower and upper fences represent the 25th and 75th percentile, and the line in between is the median. Whiskers represent the 10th and 90th percentile.

Adapter dimers are adapter sequences that are joined together and able to bind and cluster on the flow cell and generate sequencing data. Adapter dimer might cause a detrimental effect on sequencing efficiency (Maqueda, 2023). According to the QIAGEN kit handbook (Rev 03/2024), the acceptable amount of adapter dimer should not exceed 25% of the sequencing library amount, as measured in relative fluorescence unit (RFU) using capillary electrophoresis methods (e.g. Agilent Fragment Analyzer). The presence of adapter dimers was consistently higher than the recommended amount for standard 1 ng input across all protocols and increased in the protocol with higher PCR cycle numbers–standard 1 ng and ‘1 ng 2x diluted’ protocol (Figure 2). This observation indicates that input amount and number of PCR cycles seem to be more important in adapter dimer formation rather than adapter availability itself.

**Figure 2.**
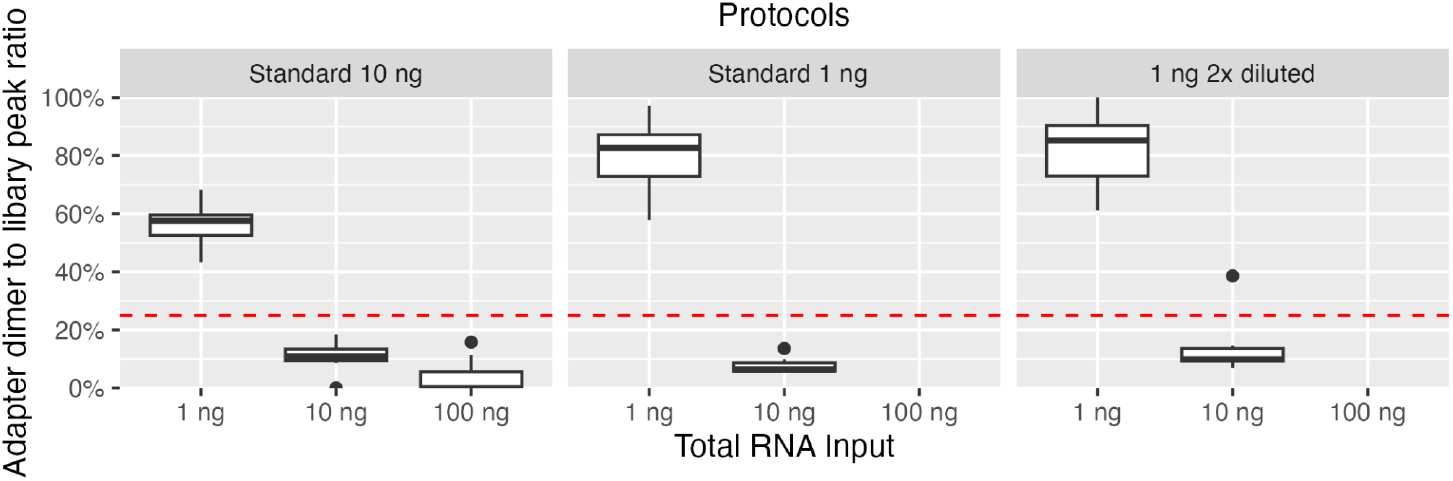
Ratio of adapter dimer to library peak (as measured by Agilent Fragment Analyzer) across different adapter dilution protocols and inputs. Lower and upper fences represent the 25th and 75th percentile, and the line in between is the median. Whiskers represent the 10th and 90th percentile. The dashed red line represents the maximum adapter dimers amount as recommended by QIAGEN.

Unique molecular identifiers (UMI) are short sequences added to DNA fragments during the preparation of sequencing libraries to identify the input DNA molecule. UMIs are added before PCR amplification and can be used to reduce errors and bias introduced by PCR amplification (König *et. al*., 2010). When preparing a sequencing library from an ultra-low input sample, PCR duplicates are often formed due to stochastic effect, GC content, and fragment length bias. However removing these duplicates while preserving true biological duplicates is possible with UMI (Parekh *et. al*., 2016).

The reads generated in sequencing prior to any quality filtering are commonly referred to as ‘raw reads’. A large proportion (∼67 – 96%) of the raw reads (before trimming by FastQC) generated across different adapter dilution protocols and inputs are PCR duplicates as it was removed after deduplication with UMI (Figure 3). The number of duplicates seems to decrease with higher input. This is evident from the higher number of raw reads removed after deduplication in the 10 ng and 1 ng input compared to the 100 ng input (Figure 3B, standard 10 ng protocol). The number of PCR cycles and adapter dilution also had a negative impact on the duplication rate, as evident from the higher number of removed reads in the ‘1 ng 2x diluted’ protocol compared to the standard 10 ng protocol.

**Figure 3.**
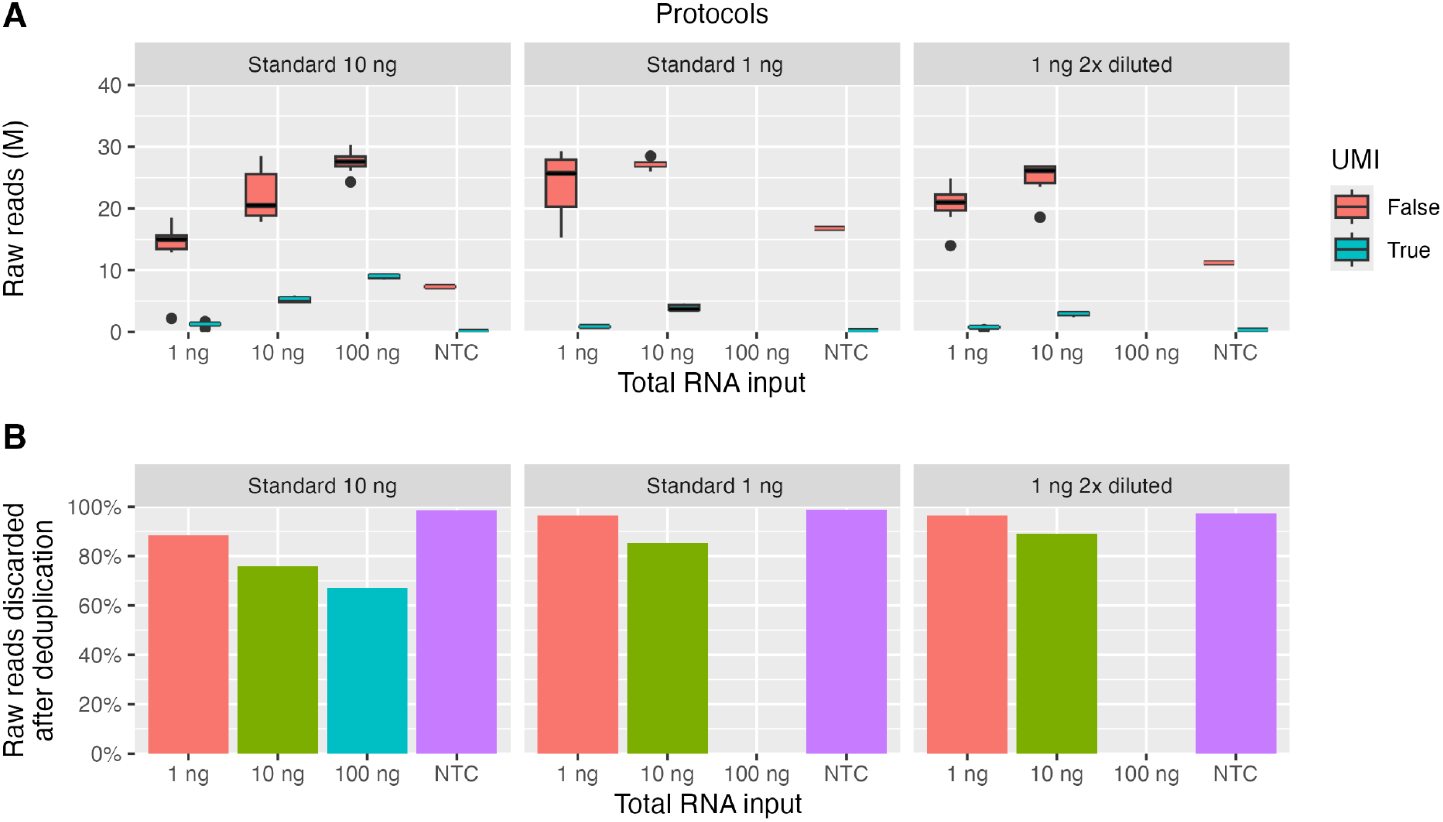
Total number of raw reads (in million) generated across different adapter dilution protocols and inputs (A). Lower and upper fences represent the 25th and 75th percentile, and the line in between is the median. Whiskers represent the 10th and 90th percentile. Average proportion of reads discarded after deduplication (B). Bar graph showing the percentage of raw reads removed after deduplication by UMI. Different colours denote different Total RNA input amounts.

To elucidate more about the severity of the duplication problem, we also analysed the number of reads that passed quality control. The filtering was done with bioinformatic tool, fastp (included in the nf-core/smrnaseq pipeline) which uses the Q-score, read length, and base complexity as criteria. Almost all reads that were retained after deduplication are high-quality reads (Figure 4). This means that UMIs help to remove low-quality reads that are byproducts of the PCR amplification.

**Figure 4.**
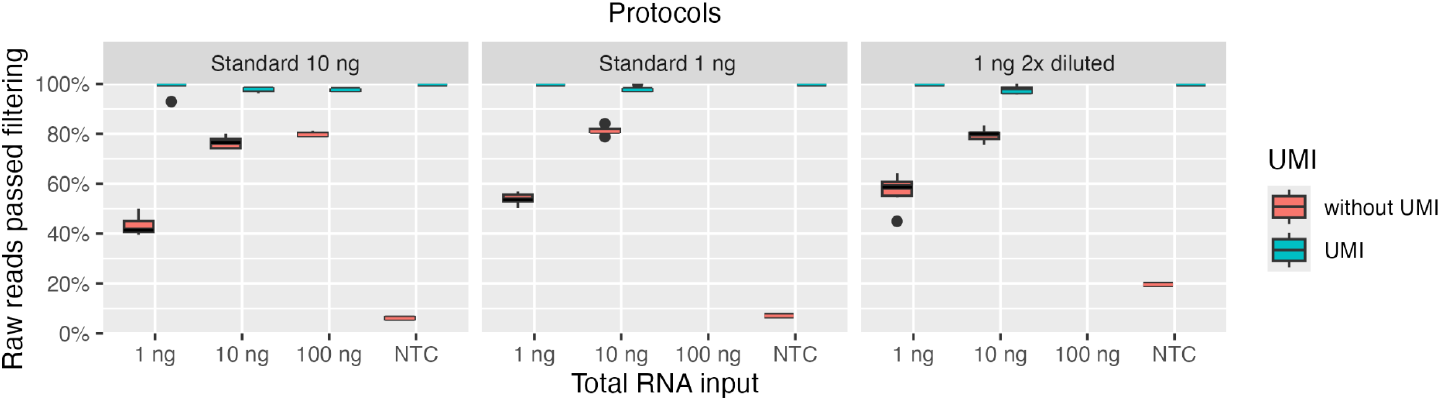
Total number of reads that passed filtering across different adapter dilution protocols and inputs. Lower and upper fences represent the 25th and 75th percentile, and the line in between is the median. Whiskers represent the 10th and 90th percentile.

To elucidate further the differences between each sample a principal component analysis (PCA) was performed based on the above parameters. The PCA plot (Figure 5) reveals distinct clustering patterns based on both the input material amount and the usage of UMIs in analysis. Along PC1, samples clearly separate according to the amount of input RNA, with larger input amounts (100 ng and 10 ng) exhibiting positive PC1 values and the smaller input amount (1 ng) showing negative PC1 values. The variation of 1 ng input amount in the ‘1 ng 2x diluted’ protocol across the PC1 indicates higher stochasticity, probably due to higher adapter dilution and higher number of PCR cycles. Furthermore, within each input amount group, samples are separated along PC2 according to whether they were analysed with or without UMIs. Notably, the 1 ng input samples show the most pronounced separation based on UMI presence and lower input amounts. This observation suggests that UMIs may be particularly critical for accurate quantification when working with limited starting material, potentially by mitigating amplification bias.

**Figure 5.**
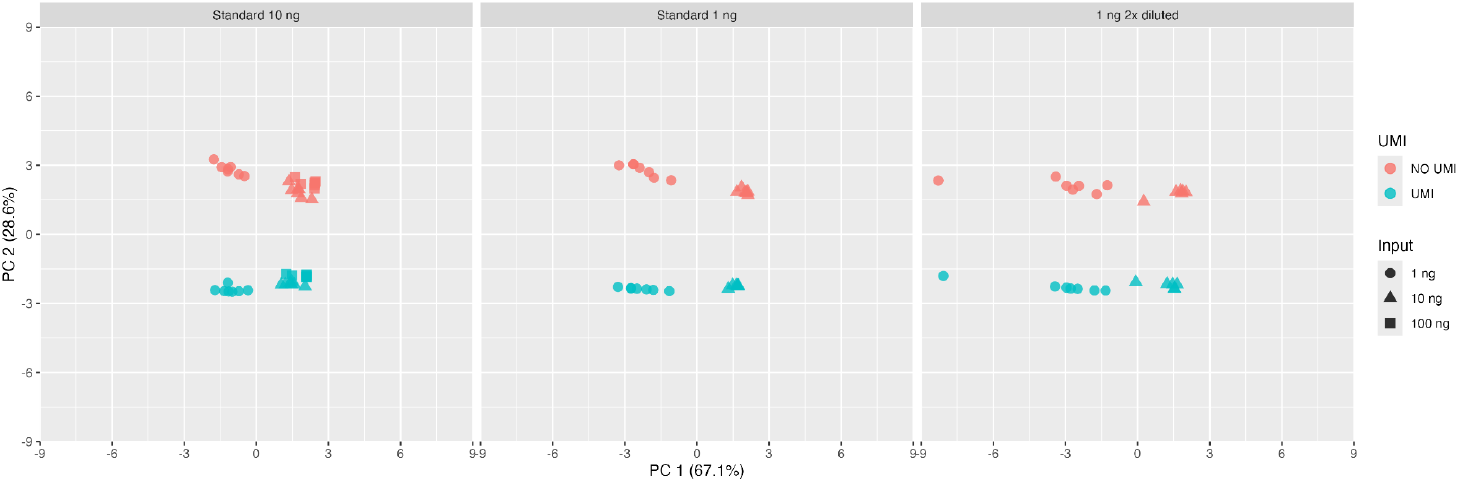
Variations in sequencing libraries across different adapter dilution protocols and inputs as visualised by principal component analysis (PCA) . Each point represents different samples, different colours denote usage of UMI and different shapes denote different input amounts.

### Library complexity

Library complexity denotes the number of true unique sequences–in this case, miRNA sequences, found in the randomly sampled subset sequencing library by miRTrace (Kang, 2018). The library complexity was affected negatively by both the total RNA Input amount and adapter dilution (Figure 6). The effect of total RNA input amount on complexity seems to be stronger compared to the effect of adapter dilution. Yet, there was little to no effect of increasing the total RNA input amount (from 10 ng to 100 ng) to complexity. On the other hand, the effect of total RNA input on complexity was not as noticeable when analysing with UMI deduplication (Figure 6B). The negative effect of adapter dilution on library complexity can be seen when comparing the 10 ng input across all protocols, both when analysing with and without UMI deduplication. It seems that, when performing deduplication with UMI, a sample does not need more than 5M reads unless the input is 100 ng or more.

**Figure 6.**
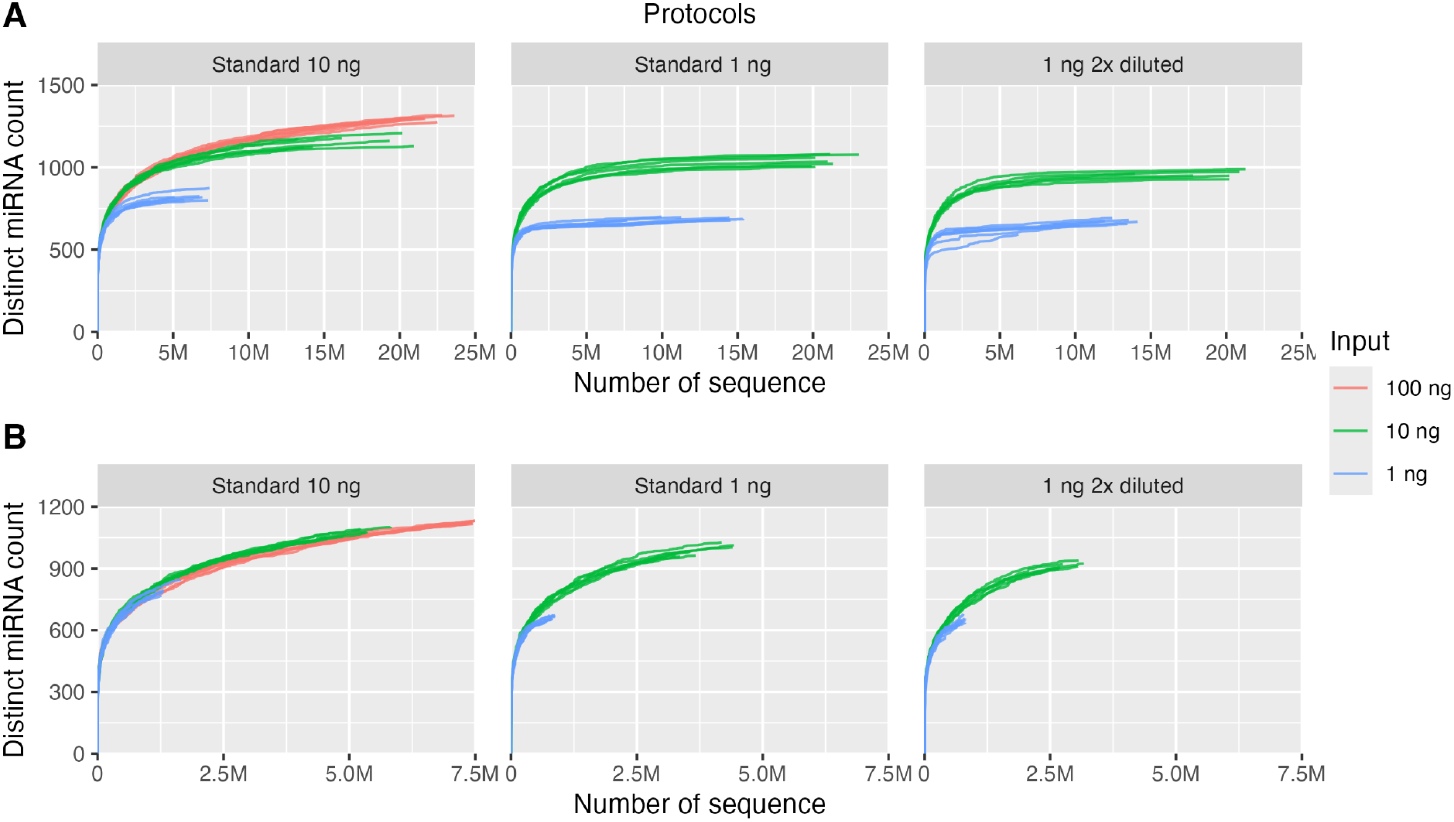
Impact of input amount on library complexity. Line plot showing the number of distinct miRNA across different adapter dilution protocols without UMI (A) and with UMI deduplication (B). Different line colours denote different input amounts.

## Conclusions

Based on this study, we concluded that the total RNA input amount and number of PCR cycles are more relevant in adapter dimer formation rather than the availability of the adapters itself. We suggest that researchers who are preparing miRNA sequencing libraries with QIAseq miRNA Library Kit should aim for the highest input possible–thus also using the lowest number of PCR cycles possible. On top of that, deduplication with UMI to remove amplification duplicates will be beneficial as it might constitute a large part of the raw reads produced during the sequencing.

We also suggest a strategy to choose the appropriate protocol when setting up a project for sequencing library preparation, where there is a large variation in the concentration of the samples, and tailoring the method to each individual sample does not seem feasible:

1. *Identify the highest possible input amount for the majority of the samples*.
2. *Adjust the input amount so it falls within a range from the common input to ten times its value*.
3. *Execute the protocol according to the input amount for the highest concentration*.

For example,

1. *For the majority of the samples the input is 1 ng and some samples can be up to 10 ng*.
2. *Prepare a workset so that the input is 1-10 ng. Aiming the highest possible amount within 5 µl (volume restriction of the kit)*.
3. *Run the 10 ng protocol*.

Even though the strategy may help with practicalities, this approach has its shortcomings. We see that having higher inputs run with low inputs protocols might reduce the library complexity and reduce sensitivity for low-expressed miRNAs. Despite that, we think that this shortcoming can be remedied by deduplication with UMI.

## Acknowledgments

We would like to thank Juliana Nardelli, Samuel J. Rulli and Paula Luu from QIAGEN for technical advice and support during this experiment.

